# Co-evolutionary analysis suggests a role for TLR9 in papillomavirus restriction

**DOI:** 10.1101/2021.04.17.440006

**Authors:** Kelly King, Brendan B. Larsen, Sophie Gryseels, Cécile Richet, Simona Kraberger, Robert Jackson, Michael Worobey, Joseph S. Harrison, Arvind Varsani, Koenraad Van Doorslaer

**Affiliations:** School of Animal and Comparative Biomedical Sciences, University of Arizona, Tucson, AZ 85721, USA; Department of Ecology and Evolutionary Biology, University of Arizona, Tucson, Arizona 85721, USA; Department of Microbiology, Immunology and Transplantation, Rega Institute, KU Leuven, 3000 Leuven, Belgium; Department of Biology, University of Antwerp, 2000 Antwerp, Belgium; The Biodesign Center for Fundamental and Applied Microbiomics, Center for Evolution and Medicine, School of Life Sciences, Arizona State University, Tempe, AZ 85287-5001, USA; The BIO5 Institute, University of Arizona, Tucson, Arizona 85721, USA; Department of Chemistry, University of the Pacific, Stockton, California, USA; Structural Biology Research Unit, Department of Integrative Biomedical Sciences, University of Cape Town, Observatory, Cape Town 7701, South Africa; Department of Immunobiology; Cancer Biology Graduate Interdisciplinary Program; UA Cancer Center, University of Arizona Tucson, AZ 85724, USA

**Keywords:** *Papillomaviridae*, Yinpterochiroptera, Yangochiroptera, Mexican free-tailed bat, innate immunity, TLR9

## Abstract

Upon infection, DNA viruses can be sensed by pattern recognition receptors (PRRs) leading to the activation of type I and III interferons, aimed at blocking infection. Therefore, viruses must inhibit these signaling pathways, avoid being detected, or both. Papillomavirus virions are trafficked from early endosomes to the Golgi apparatus and wait for the onset of mitosis to complete nuclear entry. This unique subcellular trafficking strategy avoids detection by cytoplasmic PRRs, a property that may contribute to establishment of infection. However, as the capsid uncoats within acidic endosomal compartments, the viral DNA may be exposed to detection by toll-like receptor (TLR) 9. In this study we characterize two new papillomaviruses from bats and use molecular archeology to demonstrate that their genomes altered their nucleotide composition to avoid detection by TLR9, providing evidence that TLR9 acts as a PRR during papillomavirus infection. Furthermore, we demonstrate that TLR9, like other components of the innate immune system, is under evolutionary selection in bats, providing the first direct evidence for co-evolution between papillomaviruses and their hosts.

## B. Introduction

Papillomaviruses (PVs) are circular double-stranded DNA viruses found in an extensive repertoire of hosts, including mammals, reptiles, birds, and fish (Van Doorslaer 2013; Van Doorslaer et al. 2013; 2017; 2018). In humans, roughly 400 genetically diverse papillomavirus types have been described. While a subset of these viruses is associated with (malignant) tumors, most viral types do not cause disease in immunocompetent hosts. As with humans, hosts that have been thoroughly sampled are infected with an extensive repertoire of highly diverse yet species-specific viruses. Co-evolution of virus and host alone is insufficient to explain the phylogeny of viruses in the family *Papillomaviridae*. For example, papillomaviruses infecting humans do not form a monophyletic group, within the papillomavirus family member phylogenetic tree suggests multiple evolutionary mechanisms associated with host cellular interactions and immune evasion as important factors throughout viral genome evolution (Van Doorslaer 2013; Willemsen and Bravo 2019).

Furthermore, host or tissue tropism is likely a significant determinant of host-pathogen interactions (Carey et al. 2019; Sawyer, Emerman, and Malik 2004; Taubenberger and Kash 2010). Therefore, genomic analyses that consider essential mechanisms of the viral life cycle and evolutionary pressures related to host-parasite interactions may provide a novel perspective into papillomavirus genome evolution. A broader description of animal viruses will continue to inform these efforts.

A successful infection requires that the papillomavirus DNA is delivered to the host cell nucleus. Papillomaviruses access mitotically active basal cells through lesions in the stratified epithelia of cutaneous or mucosal tissues. Following binding to cellular receptors and priming by kallikrein-8 and furin cleavage, the virus is endocytosed (Aksoy, Gottschalk, and Meneses 2017; Day and Schelhaas 2014; DiGiuseppe, Bienkowska-Haba, Guion, and Sapp 2017; Richards et al. 2006; Cerqueira et al. 2015; Day et al. 2013; Schelhaas et al. 2012). The viral DNA is transported to the Golgi before the mitosis-dependent nuclear accumulation of L2 and viral DNA near PML bodies (Day et al. 2013; Lipovsky et al. 2013; Popa et al. 2015; Aydin et al. 2017; 2014; Calton et al. 2017; DiGiuseppe, Bienkowska-Haba, Guion, Keiffer, et al. 2017; Stepp et al. 2017; Day et al. 2004).

Host cells detect a variety of viral pathogen-associated molecular patterns (PAMPs) by pattern-recognition receptors (PRRs), resulting in induction of interferon (IFN) and a potent antiviral response (Medzhitov 2007). The concerted actions of PRR signaling, specific viral-restriction factors, and viral evasion strategies determine the eventual outcome of viral infection (Bowie and Unterholzner 2008).

The papillomaviral structural proteins (L1 and L2) have no known enzymatic activity to directly counteract antiviral responses (Buck, Day, and Trus 2013; J. W. Wang and Roden 2013). Instead, we demonstrated that papillomaviruses evolved an elaborate trafficking mechanism to evade PRR sensing pathways within the cytosol (Uhlorn, Gamez, et al. 2020; Campos 2017; Uhlorn, Jackson, et al. 2020). Furthermore, millions of years of virus-host co-speciation left historical evidence of immune evasion events in these viruses’ genomes (Sorouri et al. 2020). For example, APOBEC3 has been demonstrated to restrict infection with HPV (Warren, Xu, et al. 2015; Warren, Van Doorslaer, et al. 2015). We previously demonstrated that alphapapillomaviruses are significantly depleted of TpC dinucleotides, the target for APOBEC3 mediated mutagenesis. This TpC depletion evolved as a mechanism to evade APOBEC3 mediated mutagenesis. Specifically, this depletion of the TpC content is more pronounced in mucosal alphapapillomaviruses and is correlated with significantly higher expression levels of APOBEC3 in mucosal tissues (Warren, Van Doorslaer, et al. 2015). These findings illustrate that host antiviral activity plays a critical role in regulating papillomavirus evolution and that “molecular archeology” can be used to identify these events.

Toll-like receptors (TLR) survey the extracellular and endosomal compartments and represent the first defense line against foreign invaders. TLR2 and TLR4 recognize viral glycoproteins (Blanco et al. 2010; Boehme, Guerrero, and Compton 2006; Jude et al. 2003; Bieback et al. 2002; Murawski et al. 2009; Rassa et al. 2002; M. R. Thompson et al. 2011); TLR3 recognizes double-stranded RNA (Alexopoulou et al. 2001; Bell et al. 2006; Gowen et al. 2006; Oshiumi et al. 2011; F. Weber et al. 2006; Choe 2005), TLR7 and TLR8 recognize viral single-stranded RNA (Akira, Uematsu, and Takeuchi 2006; Diebold 2004; Hemmi et al. 2002; Kawai and Akira 2006; Zucchini et al. 2008; Jurk et al. 2002; Heil 2004). The endosomal TLR9 detects unmethylated CpG motifs found in dsDNA (viral) genomes (Bowie and Unterholzner 2008; Gupta et al. 2015; M. R. Thompson et al. 2011; J. Thompson and Iwasaki 2008).

As described above, HPV particles are endocytosed (Campos 2017; Calton et al. 2017), and viral DNA could be recognized by endosomal TLR9 resulting in a downstream inflammatory immune response. We hypothesize that TLR9 may detect papillomavirus dsDNA leading to CpG depletion, similar to what we observed for APOBEC3 target motifs. Indeed, papillomavirus genomes have reduced CpG content (Warren, Van Doorslaer, et al. 2015; Upadhyay and Vivekanandan 2015). This overall dinucleotide depletion confounds the ability to demonstrate the cause of this depletion.

Bats serve as reservoirs for many viruses and and have served as the source of well-documented cross-species transmission events of pathogens responsible for a myriad of epidemics, the most notable being severe acute respiratory syndrome coronavirus 1, middle east respiratory syndrome coronavirus, Ebola virus, Marburg virus, and most recently, severe acute respiratory syndrome coronavirus 2 (Banerjee et al. 2020; Brook and Dobson 2015; Wacharapluesadee et al. 2021). It has been proposed that bats avoid immunopathological outcomes by not fully clearing a viral infection (O’Shea et al. 2014). Indeed, bats have been reported to exhibit a ‘dampened’ immune response to viral infections (Banerjee et al. 2020; Gorbunova, Seluanov, and Kennedy 2020; Xie et al. 2018; Zhang et al. 2013; Subudhi, Rapin, and Misra 2019). The complex suppression of immune response pathways is variable between several bat species (order Chiroptera) (Zhang et al. 2013; Jiang et al. 2017; Escalera-Zamudio et al. 2015; Hawkins et al. 2019). Importantly, it was demonstrated that residues involved in the ligand-binding region of the bat TLR9 protein are evolving under positive selection (Escalera-Zamudio et al. 2015; Jiang et al. 2017). This evolutionary selection of TLR9 has been proposed to contribute to the high tolerance for viral infections observed in bats (Banerjee et al. 2020; Hawkins et al. 2019). Notably, the theory of co-evolution suggests that viruses need to counter these changes in TLR9, and we should be able to detect these host-parasite interactions (Tan et al. 2017; Warren, Van Doorslaer, et al. 2015). To address this question, we determined the genomes of two novel bat papillomaviruses from *Tadarida brasiliensis* (TbraPV2, TbraPV3). Taxonomically, bats are classified into two suborders; Yinpterochiroptera and Yangochiroptera. By comparing the genomes of papillomaviruses associated with bats from either suborder, we demonstrate that TLR9 target motifs are significantly depleted and impact papillomavirus evolution. Furthermore, we extend existing data showing that Yangochiroptera TLR9 is evolving under diversifying selection. This argues that papillomavirus genomes are evolving in response to a host change. This is the first direct evidence of PVs evolving in response to host evolutionary changes, thus providing direct evidence for co-evolution. Also, these data argue that TLR9 is a restriction factor for papillomavirus infection.

## C. Results

### C1. Sampling, sample processing and viral metagenomics

We identified two circular contigs (circular based on terminal redundancy) that had similarities to papillomavirus sequences. We mapped the raw reads to the assembled genomes using BBmap (Bushnell 2014)to determine the read depth. For both the genomes we had a 22-25X coverage depth across the whole genome with 1200-1300 mapped reads.

### C2. New bat associated papillomaviruses cluster with previously identified Chiropteran viruses

Using a metagenomics approach, we determined the genomes of two novel circular dsDNA viruses. The open reading frames of these putative new viruses were determined using PuMA (Pace et al., 2020). This analysis identified the typical papillomavirus open reading frames (E6, E7, E1, E2, L1, and L2) and the spliced E1^E4 and E8^E2 mRNAs), suggesting that we recovered the genomes of two papillomaviruses associated with Mexican free-tailed bats (*Tadarida brasiliensis*). The current papillomavirus taxonomy is based on sequence identity across the L1 open reading frame. If two viruses share more than 60%, they fall into the same genus. Species within a genus group viral ‘types’ that share between 70 and 90% sequence identity. A new papillomavirus type shares less than 90% sequence identity with other viruses (Van Doorslaer et al. 2018; Bernard et al. 2010; de Villiers et al. 2004). Both identified viruses share less than 90% identity with their closest relatives (**Figure 1B**). In consultation with the international animal papillomavirus reference center (Van Doorslaer and Dillner 2019), we name these two novel papillomaviruses TbraPV2 (*GenBank #* MW922427) and TbraPV3 (*GenBank #* MW922428), respectively. TbraPV2 is 8093 bp long, while TbraPV3 is 8037 bp long. Based on pairwise sequence identity in the L1 open reading frame, both viruses are most closely related to TbraPV1 (**Figure 1B**). TbraPV2 shares 81.7% sequence identity with TbraPV1 and likely represents a new species in this as of yet unclassified genus. TbraPV3 shares 60.4% identity with TbraPV1, placing it in the same genus. However, the phylogenetic tree shown in **Figure 1** places TbraPV3 as an outgroup to a clade that contains HPV41, EdPV1, TbraPV1, TbraPV2. Therefore, the evolutionary history of these viruses does not support the current L1-based taxonomy.

**Figure 1.**
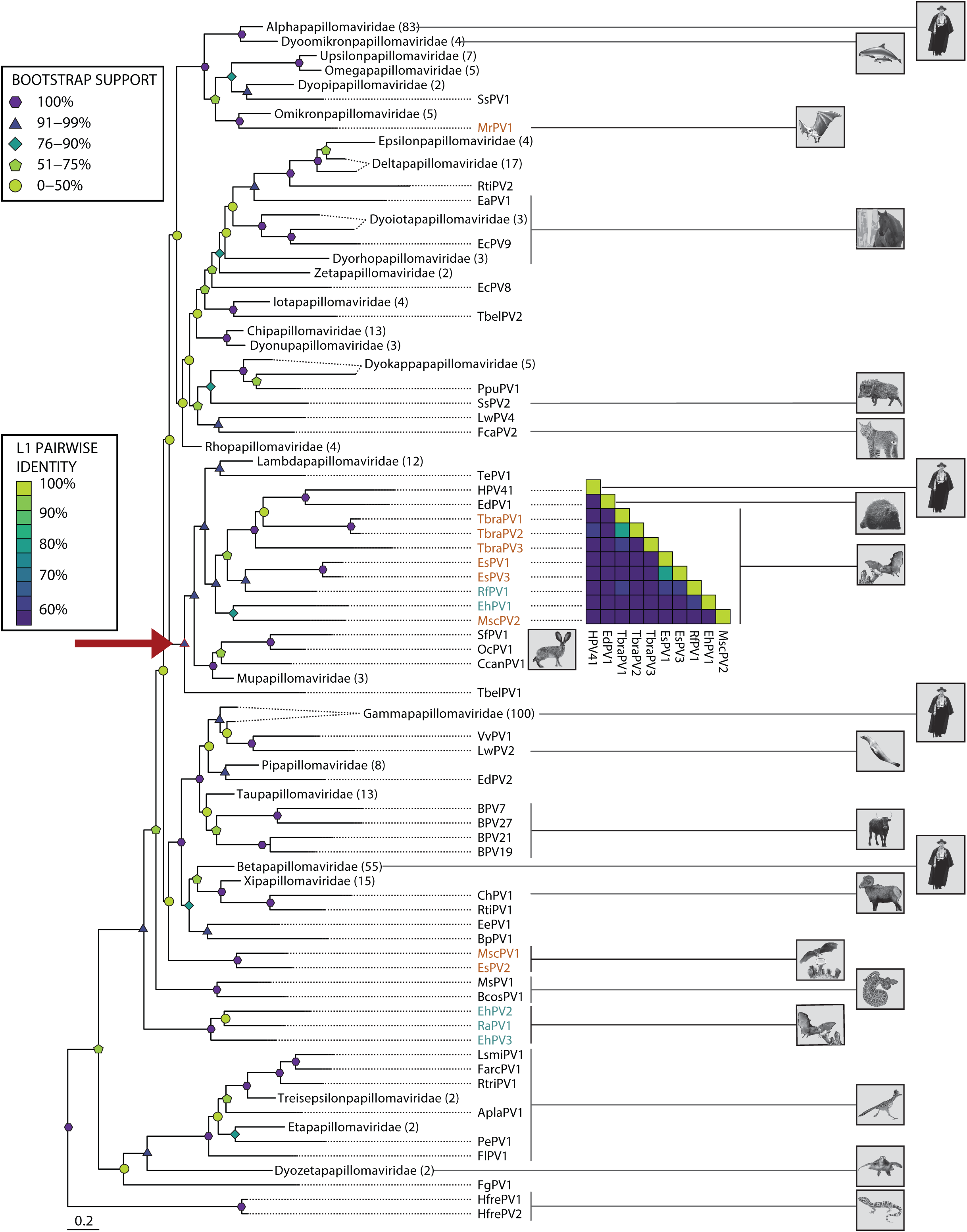
Evolutionary relationship of novel bat papillomaviruses. (A) Maximum-likelihood phylogenetic tree inferred using concatenated E1, E2, and L1 protein sequences. Papillomaviruses associated with *Chiroptera* are highlighted Yangochiroptera (orange) and Yinpterochiroptera (red). Papillomavirus genera are collapsed (number of types within each genus are indicated in parentheses). Bootstrap generated branch support values are given using symbols and color gradient. Host species are indicated using Sonoran Desert dwelling animals. The red arrow indicates the subtree used for further analyses throughout the manuscript. (B) Pairwise identity plot with percentage pairwise identities provided in colored boxes for the L1 nucleotide sequences.

It has been demonstrated that co-speciation between PVs and their hosts is a major contributor to the papillomavirus’s evolutionary history (Van Doorslaer 2013; Gottschling et al. 2007; 2011). In support of this notion, these novel bat papillomaviruses cluster with other previously described Chiropteran papillomaviruses (**Figure 1A**). However, as for other papillomavirus-host relationships, bat-associated viruses are paraphyletic, suggesting that other evolutionary mechanisms like intra-host divergence or niche adaptation likely contribute to the papillomavirus phylogenetic tree (Buck et al. 2016; Van Doorslaer 2013).

### C3. Chiropteran PVs co-speciated with their hosts

While TbraPV2 and TbraPV3 cluster together with several other Chiropteran viruses, the larger clade consists of viruses infecting a wide array of mammals (red arrow in **Figure 1A**). In addition to 6 species of Chiroptera, the subtree contains 16 host species classified in 5 mammalian orders. We wanted to compare the evolutionary history of these diverse viruses to their hosts. Due to intra-host divergence and niche adaptation, papillomaviruses infecting the same host can be found in multiple phylogenetic tree clades. To ensure that viruses with a similar tissue tropism and evolutionary history are compared, we extracted a subtree from the maximum likelihood tree (**Figure 2**) (Smeele et al. 2018). This clade contains the newly identified TbraPV2 and TbraPV3 embedded within the largest monophyletic Chiroptera papillomavirus clade (red arrow in **Figure 1A**). We used a tanglegram to address our hypothesis of virus-host co-evolution (**Figure 2A**). In this analysis, nodes in the host and virus phylogeny are rotated to optimize tip matching. Similarities between the host and virus phylogenetic relationships are indicated by parallel lines linking the virus to its host in their respective trees. Conversely, mismatches in the evolutionary history of the host and the virus show overlapping connecting lines. While there are some overlapping connections between papillomaviruses and their hosts, most virus-host pairs support the idea of co-speciation.

**Figure 2.**
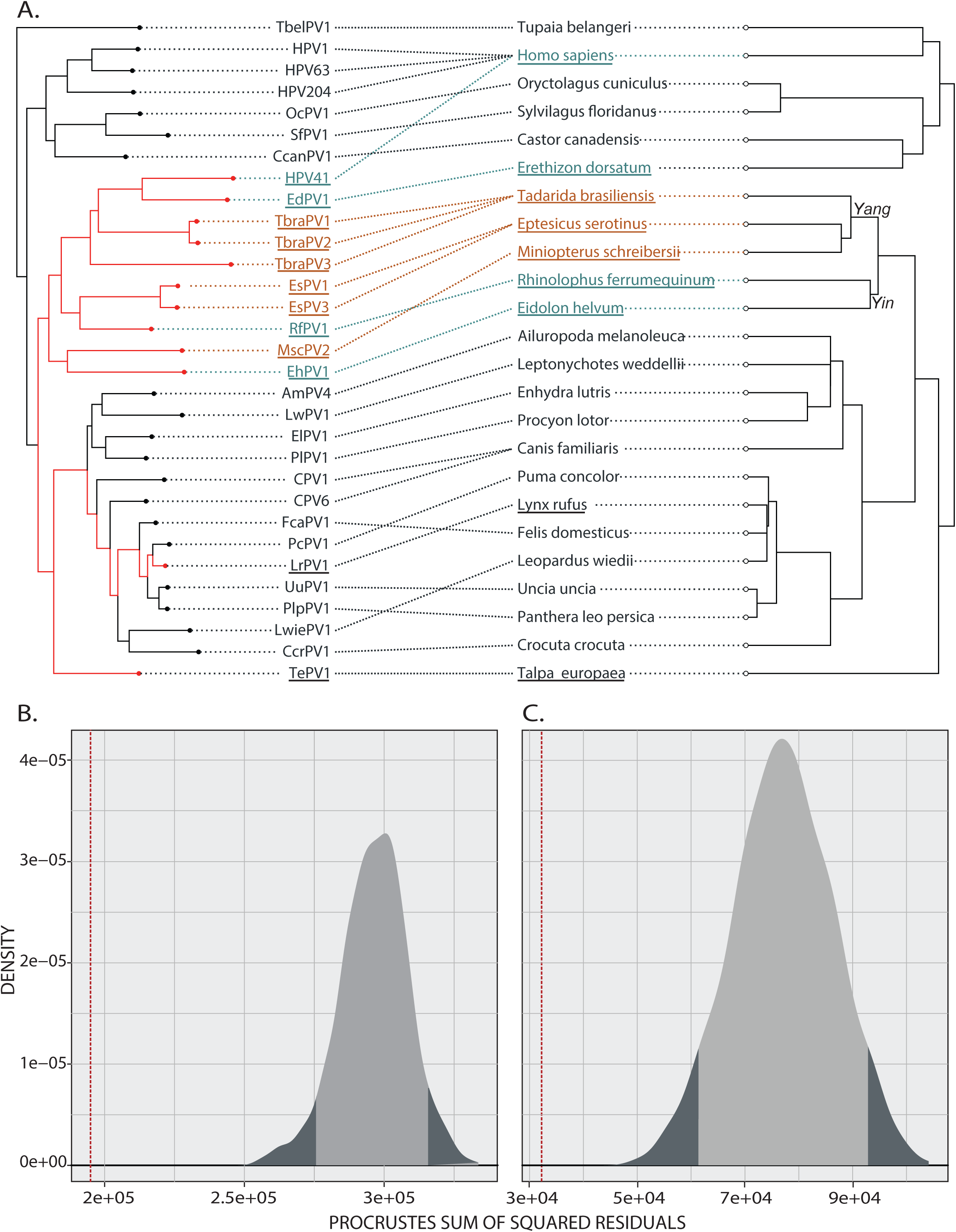
Co-evolution of papillomaviruses. (A) Optimized tanglegram between subtree based on concatenated E1–E2–L1 maximum-likelihood phylogenetic tree (see Figure 1) and associated host species. Host species tree was downloaded from www.timetree.org. Papillomaviruses are linked to their host phylogenies. Papillomaviruses associated with Chiroptera are highlighted Yangochiroptera (orange) and Yinpterochiroptera (blue). (B) Procrustean Approach to Cophylogeny analysis based on the interaction network and phylogenies shown in (A) supports that papillomaviruses coevolved with their hosts. The observed best-fit Procrustean super-imposition (red dotted line line) lies outside of the 95% confidence interval (shaded area of the curves) of the distribution of network randomizations in the null model. (C) As in (B) but using a subset of the interaction network and phylogenies (indicated by red arrow in (A))

To formally quantify the degree of co-speciation, we focused on two datasets. First, we used the phylogenetic tree shown in **Figure 2**. Because it was previously shown that members of the *Lambdapapillomavirus* genus co-evolve with their hosts (Rector et al. 2007), we also tested a smaller subtree to avoid skewing the results (indicated with the red arrow in **Figure 2**). The host and virus phylogenetic trees were compared using the Wasserstein distance, estimated to be 0.205 and 0.284 for the larger and smaller datasets, respectively. A Wasserstein distance of 0 indicates that both trees are topologically identical, while a value of 1 indicates complete lack of congruence between both trees (Lewitus and Morlon 2016). Therefore, the host tree predicts the virus tree, suggesting a role for co-speciation.

Also, we used the Procrustean Approach to Cophylogeny (PACo). This approach evaluates congruency between distance matrices for each virus and associated host phylogenies (Balbuena, Míguez-Lozano, and Blasco-Costa 2013; Hutchinson et al. 2017). The observed best-fit Procrustean super-imposition (1.08E5 and 3.22E4 for the larger and smaller dataset, respectively; denoted by the red dotted line) lies outside of the 95% confidence interval of the ensemble of 1000 network randomizations in the null model (**Figures 2B and C**). Therefore, the data allow us to reject the null hypotheses that the papillomavirus host tree does not predict the virus tree and supports co-speciation as an important process for the evolution of this subclade of PVs and their hosts (Balbuena, Míguez-Lozano, and Blasco-Costa 2013; Hutchinson et al. 2017).

### C4. Yangochiropteran viruses have a reduced CpG content

We previously demonstrated that millions of years of virus-host co-speciation left historical evidence of this virus-host arms-race in the papillomavirus’ genomes. For example, the mucosal, cancer-causing alphapapillomaviruses have a reduced TpC dinucleotide content, presumably due to evolutionary adaptations to APOBEC3 editing (Warren, Van Doorslaer, et al. 2015). To extend these studies, we calculated the observed/predicted ratio for each dinucleotide in the viruses shown in **Figure 2**. A ratio close to 1 indicates that a dinucleotide is seen in the sequence as often as expected based on each sequence’s nucleotide composition. Values lower than 1 suggest that a dinucleotide is depleted. While the ApC, ApT, GpT, TpA, and TpC ratios are significantly lower than 1 (one-sample t-test p-value < 0.001), we observed the most significant decrease in the CpG dinucleotide ratio (**Figure 3**). The median CpG content for these evolutionarily related viruses is 0.46. However, we noticed that the distribution has a long tail towards even lower values, suggesting that some viruses have a further reduced CpG content (**Figure 3**).

**Figure 3.**
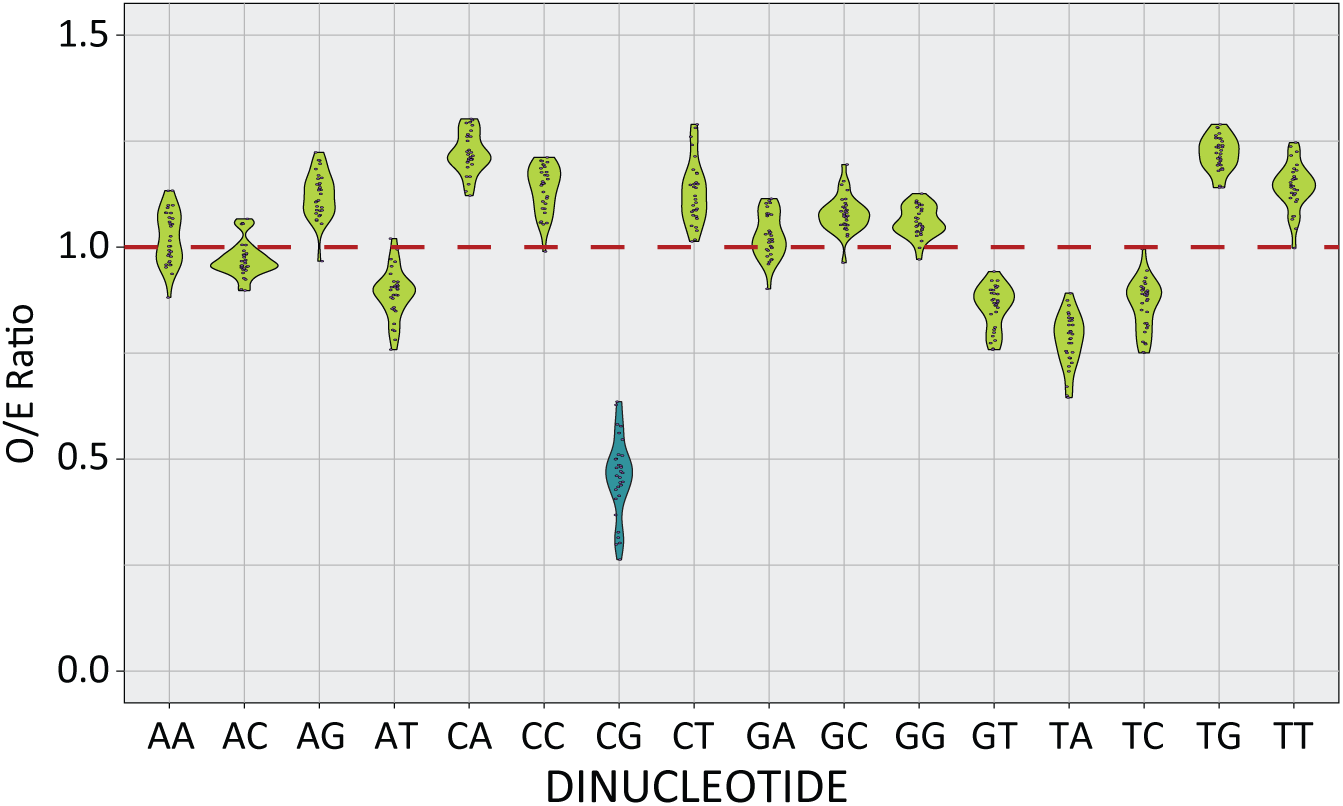
CpG dinucleotide sequences are significantly depleted in papillomavirus genomes. The observed vs. expected (O/E) ratios of each dinucleotide in the papillomavirus genomes sequences shown in Figure 2 were calculated using a custom wrapper around the CompSeq program from the EMBOSS software suite. The red line indicates that the sequence is seen as often as would be expected by chance.

When we plotted the CpG ratio for each virus on a phylogenetic tree (**Figure 4A**), it became clear that the genomes of a subset of bat-associated papillomaviruses have a further decreased CpG content. The order Chiroptera consists of two suborders, Yinpterochiroptera and Yangochiroptera (Lei and Dong 2016; Teeling et al. 2002; Springer et al. 2001). When we associated the viruses with their Yinpterochiroptera and Yangochiroptera hosts, the data demonstrates that the Yangochiropteran papillomavirus genomes have even lower CpG values (orange bars in **Figure 4A**) when compared to the Yinpterochiroptera and other related papillomaviruses in the same phylogenetic clade (blue bars) and members of a closely related clade (grey bars). Indeed, when combined, the Yangochiropteran papillomavirus genomes have a significantly reduced CpG ratio when compared to the other groups (**Figure 4B** ANOVA with posthoc Tukey test).

**Figure 4.**
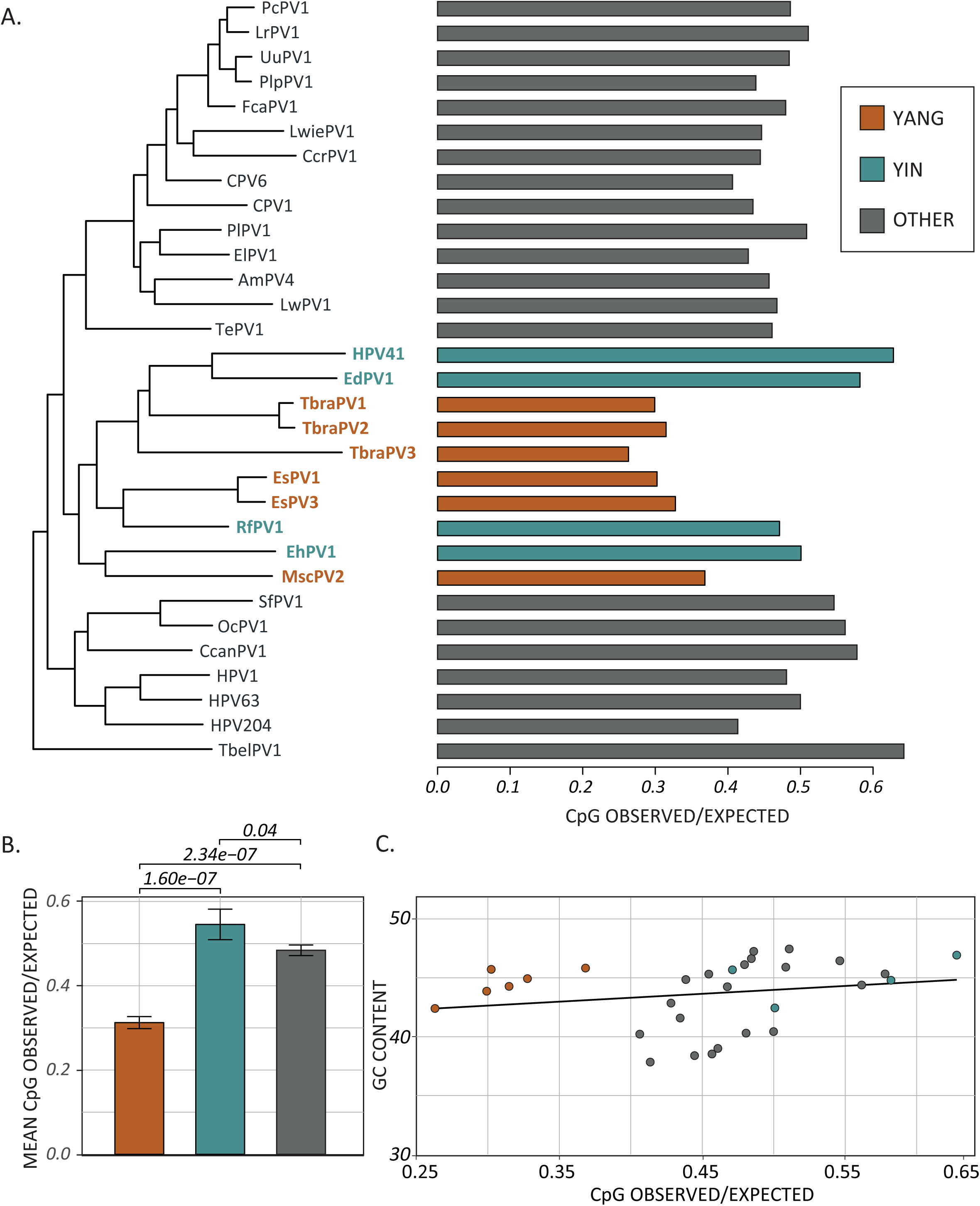
CpG content is significantly lower in papillomaviruses associated with Yangochiroptera compared to related viruses. (A) A maximum likelihood phylogenetic tree is shown comparing the O/E ratios of CpG dinucleotides. Viruses infecting Yangochiroptera (red), Yinpterochiroptera (green), and related hosts (grey) are indicated. (B) Mean (+/- standard deviation) CpG observed vs. expected (O/E) ratios for each group of viruses are compared using a one-way ANOVA with Tukey’s posthoc test. (C) CpG observed vs. expected (O/E) ratios are compared to total GC content for each viral genome in (A).

This reduction in the CpG ratio could be due to an overall lower GC content. We compared the CpG ratio to total genomic GC content (**Figure 4C**). This analysis demonstrates that there is no correlation between the decreased CpG ratio and the total GC content (linear regression: R^2^ = 0.008, p-value = 0.27), and the reduced CpG ratio is not simply due to a lower GC content.

The viruses infecting bats in the Yangochiroptera suborder have a reduced CpG content, raising the possibility that the host species is influencing the CpG ratio and evolutionary trajectory of these viruses.

### C5. CpG depletion alters codon usage without changing amino acid composition

The CpG dinucleotide is present in 8 codons coding for five different amino acids. Therefore, reducing CpG dinucleotides is expected to lead to a bias in codon usage or amino acid composition. Roughly 85% of the papillomavirus genome codes for viral proteins (Van Doorslaer et al. 2013; 2017). The viral genome contains several overlapping ORFs (Van Doorslaer 2013; Van Doorslaer and McBride 2016). In some cases, this is a short overlap between the 3’ end of one ORF and the 5’ end of the downstream ORF (e.g., E6 and E7). In other cases, one ORF is wholly embedded within the coding region of another ORF (e.g., E4 embedded in E2, or E8 within E1). Since these overlapping regions are evolutionarily constrained at multiple codon positions (i.e., codon position 3 in frame 1, would be codon position 2 in the overlapping frame (Miyata and Yasunaga 1978), these overlapping regions were removed from the data (Materials and Methods).

We determined codon usage tables for the non-overlapping coding sequences for each of the papillomaviruses in the subtree described in **Figure 2**. These codon usage tables were compared using the Emboss ‘codcmp’ tool to calculate codon usage differences. The more diverse the codon usage, the larger the differences between both tables. This analysis shows that compared to each other, the Yangochiroptera papillomaviruses codon usage is more similar than when other viruses are compared (**Figure 5A**). This suggests that the reduction in CpG leads to a more restricted availability of codons. The relative synonymous codon usage (RSCU) value is the ratio of the observed frequency of one specific synonymous codon to the expected frequency (i.e., no codon usage bias). This ratio is an important measure of codon usage bias (Sharp, Tuohy, and Mosurski 1986). RSCU values higher than 1.6 and lower than 0.6 indicate overrepresented and underrepresented codons, respectively (Wong et al. 2010). CpG containing codons (underlined) are significantly underrepresented in this dataset. For most amino-acids, CpG containing codons are further reduced in the Yangochiroptera papillomaviruses. Of note, in the case of arginine, the CpG containing codons are statistically significantly depleted when compared to the related viruses (**Figure 5C**). Since these codons’ CpGs are located in the 1^st^ and 2^nd^ position of the codon, this depletion suggests that non-silent mutations are evolutionarily preferred over maintaining a relatively high CpG content. Of note, despite the biased codon usage, there is no change in the amino acid composition of the different viral proteins (**Figure 5B**). Thus, despite reducing CpG content, the Yangochiroptera papillomaviruses are likely coding for similar proteins. These observations likely explain the more restricted codon usage seen in **Figure 5A**.

**Figure 5.**
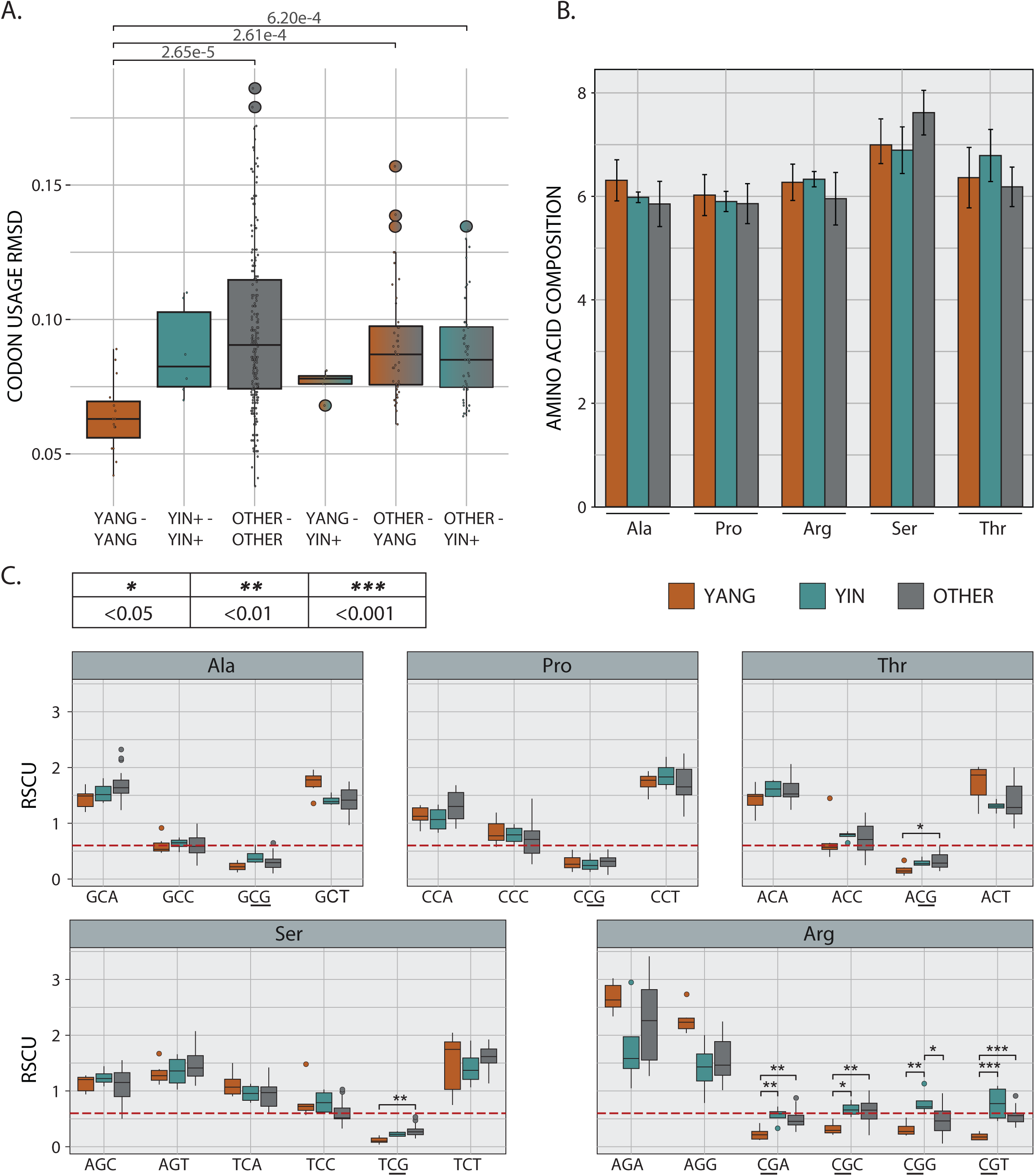
Yangochiroptera papillomaviruses have a restricted codon usage. (A) Codon usage tables for each virus in Figure 2 were compared using the ‘codcmp’ program from the EMBOSS software suite. Root-mean-square deviation (RMSD) values for each pairwise comparison are plotted as Box-and-whisker plots with the outliers (colored circles) identified using Tukey’s method. Individual values are shown as a single black dot. (B) Amino-acid composition was calculated as described in materials and methods. Mean values +/- standard deviation is plotted. (C) RSCU values for the indicated amino acid/codons were calculated and plotted as Box-and-whisker plots with the outliers (colored circles) identified using Tukey’s method. RSCU values for each amino acid were compared using a two-way ANOVA with Tukey’s posthoc test. Significance is indicated as shown in the legend.

### C6. Natural selection in the TLR9 of *Yangochiroptera* bats

Specific residues within the Chiropteran TLR9 have been demonstrated to be under positive selective pressure (Escalera-Zamudio et al. 2015; Jiang et al. 2017). To determine whether diversifying selection differentially affected Yangochiroptera and Yinpterochiroptera TLR9, we constructed a maximum likelihood phylogenetic tree. As was previously reported, bat TLR9 sequences formed a monophyletic clade separate from other eutherian sequences (data not shown), not as a sister-group to carnivores, ungulates, and cetaceans as is seen for other proteins (Tsagkogeorga et al. 2013). We used RELAX (Wertheim et al. 2015), adaptive Branch Site Relative Effect Likelihood (Smith et al. 2015), and Fixed effect Likelihood tests (Kosakovsky Pond and Frost 2005) to detect evidence for evolutionary selection (materials and methods). RELAX demonstrated that evolutionary selection intensified (K = 6.07; LR = 21.05) within the Yangochiroptera compared to the Yinpterochiroptera. Furthermore, aBSREL recovered evidence for episodic diversifying selection on the branches leading to the Yangochiroptera (**Figure 6B**). Finally, FEL identified 7 sites under diversifying selection within the Yangochiroptera.

**Figure 6.**
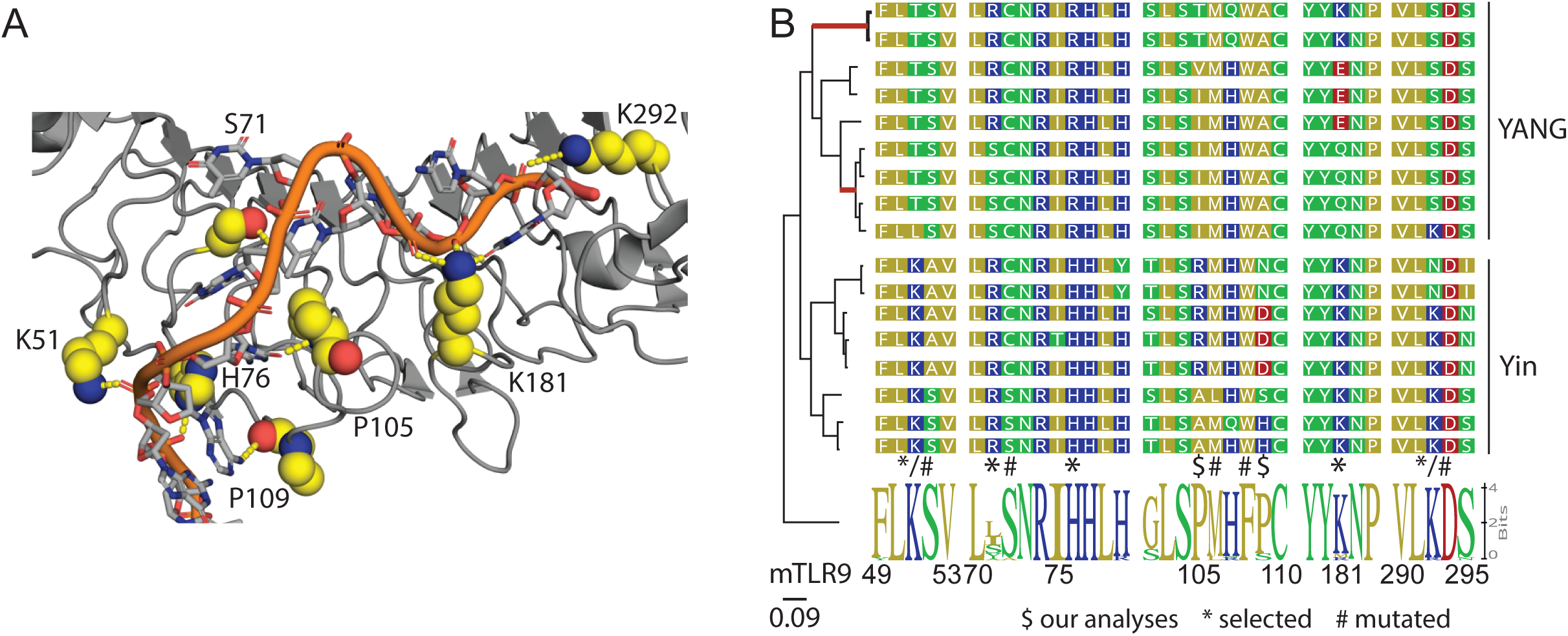
Yangochiroptera TLR9 is evolving under diversifying selection. (A) Structure of horse TLR9 in complex with agonistic DNA (PDB: 3WPC) (Ohto et al. 2015). Amino acids of interest are highlighted. (B) Maximum likelihood phylogenetic tree of mammalian TLR9 sequences clusters Yangochiroptera and Yinpterochiroptera separate from the mammalian TLR9. Red branches display evidence of episodic diversifying selection as identified by aBRSEL (Smith et al. 2015). Alignments show sequences of interest. The sequence logo is based on the alignment of 29 non-chiropteran TLR9 sequences. Numbering is based on the mouse TLR9. Residues indicated with $ were identified as being selected using FEL. Residues highlighted with * were previously identified as evolving under diversifying selection (Escalera-Zamudio et al. 2015), while residues with # were shown to be functionally important through site directed mutagenesis (Ohto et al. 2015.

We mapped a subset of these residues as well as residues previously identified to be under diversifying selection (Escalera-Zamudio et al. 2015), and sites shown to be functionally important for target recognition (Ohto et al. 2015) onto the structure of TLR9 bound to target DNA (**Figure 6A**). Many of the sites are highly variable when compared to the mammalian consensus (**Figure 6B**). Notably, there are apparent differences between the TLR9 sequence of Yangochiroptera and Yinpterochiroptera, specifically within the DNA recognition motif. *In silico* mutation analysis suggests that these Yangochiroptera specific changes would alter how TLR9 recognizes its target DNA. Overall, these mutations lead to a reduction in the positive surface charge of TLR9. Within the Yangochiroptera, K51T leads to the loss of an ionic interaction with the phosphate backbone. The Arg at position 76 is much larger than the canonical His at this position, which will presumably impact the interaction with the DNA. While the P105 residue Vanderwaals bonds with the C6 and T9 residues in the crystalized DNA, the I105 is too bulky to occupy the same conformation as the proline and will likely lead to a loss of the observed bend in the protein at this position. K181 is involved in ionic interactions with the DNA backbone. Q181 has no charge and is too short to interact with the DNA sidechain. E181 will likely charge repel the DNA backbone. Finally, K292 interacts with the DNA backbone, but this interaction in absent in Yangochiroptera TLR9 due to the Ser residue at this position. Overall, the Yangochiroptera TLR9 DNA binding domain is predicted to be functionally different from the Yinpterochiroptera and the other mammalian TLR9 molecules.

### C7. CpG depletion points toward a TLR9 signature

Our data demonstrate that the <u>Yangochiroptera</u> TLR9 protein is under selective pressure, and papillomaviruses that infect these hosts have a decreased CpG content. We hypothesized that DNA recognition by TLR9 would lead to a decrease in CpG in the context of a TLR9 specific PAMP. Thus, a specific (set of) tetramers should be depleted within the Yangochiroptera. We calculated the observed/expected ratio for all tetramers and focused on those tetramers with a central CpG (NCGN; **Figure 7A**). This initial analysis indicates that ACGT, GCGT, TCGA, and TCGT are diminished in Yangochiroptera. We calculated the average tetramer ratio for each group of viruses to normalize for the differences in overall CpG content between Yangochiroptera and other viruses, as in **Figure 4**. We compared the proportion of Yang to Yin, Yang to other, and Yin to other (**Figure 7B**) for each tetramer. For example, in Yangochiroptera papillomaviruses, ’ACGT’ is depleted three- to four-fold compared to other (brown) or Yinpterochiroptera papillomaviruses (orange bar), respectively. Conversely, this tetramer is not depleted when Yinpterochiroptera and other viruses are compared (blue bar). As expected, CpG containing tetramers are depleted in Yangochiroptera papillomaviruses. Using a bootstrap method based on 1000 randomly shuffled sequences (Materials and Methods), the ‘ACGT’ tetramer was identified as significantly depleted within Yangochiroptera specific viruses (**Figure 7C**). This tetramer is identical to the experimentally validated core mouse TLR9 recognition motif but is different from the human TLR9 PAMP (TCGT) (Pohar et al. 2015; Krieg et al. 1995; Yi et al. 1998; G. Sen et al. 2004; Hartmann and Krieg 2000). This suggests that papillomaviruses associated with Yangochiroptera specifically deplete CpG dinucleotides in the context of a known TLR9 PAMP.

**Figure 7.**
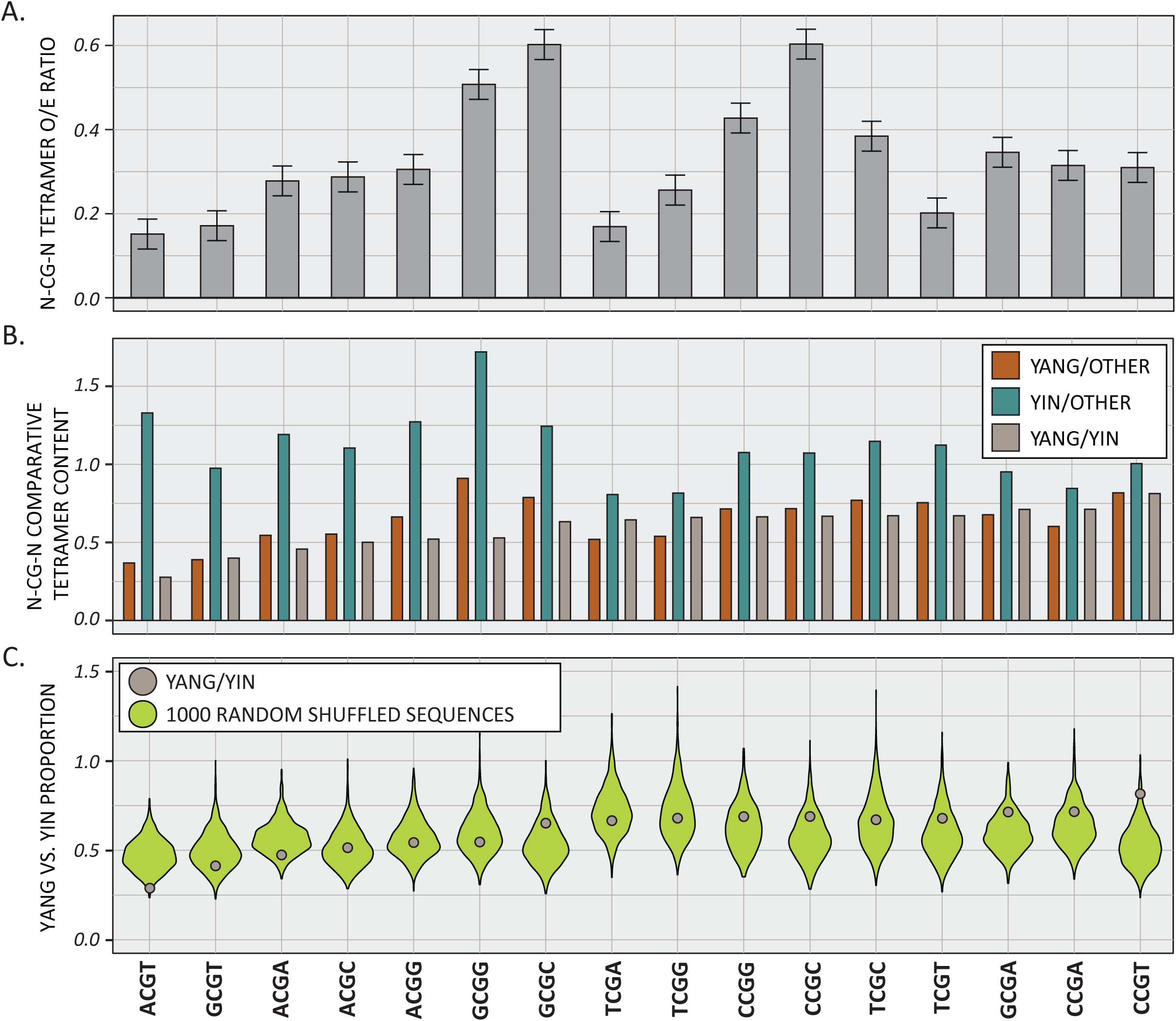
Yangochiroptera papillomaviruses depleted a TLR9 recognition motif from their genomes. (A) The observed vs. expected (O/E) ratios of each N-CG-N tetramer in the Yangochiroptera papillomavirus genomes sequences were calculated using a custom wrapper around the CompSeq program from the EMBOSS software suite. Mean values +/- standard deviation are plotted. (B) The observed vs. expected (O/E) ratios of each N-CG-N tetramer in the different groups was calculated as in A. The proportion of these ratios are shown to provide a normalized view of tetramer depletion across papillomavirus genomes shown in Figure 2. (C) The Yang vs. Yin N-CG-N proportion (as in B) are plotted as brown dots and compared to 1000 randomly shuffled sequences (green violin) plots. Only ACGT is statistically underrepresented in the Yangochiroptera.

## D. Discussion

The data presented here advance our understanding of papillomavirus evolution and host-pathogen interactions. Specifically, we provide evidence that papillomavirus genomes have evolved to avoid detection by TLR9. The implications of this finding for papillomavirus biology are discussed below.

### D1. Viruses infecting bats in the suborder Yangochiroptera deplete CpG in a TLR9 dependent manner

We demonstrate that the genomes of viruses isolated from specific bat species have a highly reduced CpG content. A significant reduction of CpG sites in papillomavirus genomes has been previously documented (Warren, Van Doorslaer, et al. 2015; Upadhyay and Vivekanandan 2015). However, the reason for this depletion is unclear. Of note, in mammalian genomes, CpGs are rare outside of so-called CpG islands (Illingworth and Bird 2009). This is believed to be mainly due to the observation that methylated CpGs are prone to deamination, resulting in C → T mutations, leading to a depletion of CpG sites in the mammalian genomes over evolutionary time.

Our data in **Figure 3** show an increase in TpG and CpA. However, this increase does not appear to be of the same magnitude as the dramatic reduction in CpG seen in the same dataset. The zinc-finger antiviral protein (ZAP) acts as a broad-spectrum antiviral restriction protein that recognizes CpG rich viral RNA, leading to RNA degradation and inhibition of translation (Gao 2002). Interestingly, it appears that ZAP exploits host CpG suppression to identify non-self RNA. This may explain why multiple RNA viruses have reduced CpG content (Cheng et al. 2013; Greenbaum et al. 2008), independently from CpG methylation as described above (Takata et al. 2017). ZAP was recently shown to restrict the replication of vaccinia virus Ankara (Peng et al. 2020) and HCMV (Lin et al. 2020), demonstrating that ZAP recognizes CpG rich viral RNA and can restrict CpG rich DNA viruses. However, papillomavirus genomes are generally CpG depleted (Warren, Van Doorslaer, et al. 2015; Upadhyay and Vivekanandan 2015). Furthermore, the consensus recognition site for murine ZAP was identified as CN7GNCG. In this motif, the CG dinucleotide acts as the essential element, while the G further enhances binding affinity 10-fold (Luo et al. 2020). Our tetramer analysis does not identify a downregulation of GNCG in Yangochiroptera specific papillomaviruses (data not shown). Therefore, it appears unlikely that ZAP plays a vital role during papillomavirus infection. However, this would need to be demonstrated experimentally.

In contrast, we provide evidence that the depletion of CpG in papillomavirus genomes is, at least in part, due to the need to avoid detection by TLR9. Unmethylated CpG DNA motifs are recognized by TLR9, leading to an interferon and inflammatory cytokine-mediated antiviral response (Kawai and Akira 2006). By carefully analyzing the CpG content of related Chiropteran papillomaviruses, we demonstrate that viruses isolated from Yangochiroptera have a further decreased CpG content. Importantly, we demonstrate that the Yangochiroptera TLR9 protein is evolving under diversifying selection, specifically sites implicated in DNA recognition. Finally, by analyzing tetramer motifs, we show that Yangochiroptera are specifically depleted in ACGT, a known TLR9 recognition motif. Together these data demonstrate that Yangochiroptera papillomaviruses deplete CpG, in the context of a TLR9 recognition motif, presumably in response to evolutionary changes within the TLR9 protein. This has important implications for papillomavirus biology and evolution.

### D2. Recognition of papillomavirus DNA in the endosomes during infectious entry

Shortly after entry, papillomavirus virions are trafficked from early endosomes into acidic late endosome and multivesicular bodies, leading to capsid disassembly and uncoating viral DNA (Campos 2017). Presumably, this exposes the viral DNA to TLR9, leading to an antiviral response. Importantly, TLR9 specifically recognizes unmethylated CpG motifs. Several studies have investigated the methylome of oncogenic human papillomaviruses (Johannsen and Lambert 2013). While these studies have demonstrated that the viral DNA is methylated under specific conditions, it is unknown whether the packaged viral genome contains methylated CpG sites. However, we have some clues that would suggest that viral DNA inside the virion is likely hypomethylated. DNA methyltransferase 1 (DNMT1) is the primary cellular enzyme responsible for maintaining DNA methylation patterns after replication. The DNMT1 protein was found enriched in undifferentiated cells and is reduced as cells differentiate (G. L. Sen et al. 2010, 1). Therefore, it is likely that the reduction in DNMT1 levels will lead to a loss of methylation on the viral genomes destined for packaging and infection of the new tissue.

Differentially methylated CpG dinucleotides are present within consensus E2 binding sites in the viral upstream regulatory region (McBride 2013). The binding of E2 to these binding sites is important for viral replication, transcription, and proper partitioning of the viral genomes to daughter cells (McBride 2013). In many viruses, the full-length E2 protein either activates or represses viral transcription in a dose-dependent manner (Bouvard et al. 1994; Fujii et al. 2001; Thierry and Yaniv 1987; Steger and Corbach 1997). CpG methylation of these sites inhibits E2 binding, presumably altering E2-mediated control of E6/E7 oncogene expression (Thain et al. 1997; Vinokurova and von Knebel Doeberitz 2011). However, the impact of changes to E2BS methylation during cellular differentiation is not understood (Burley, Roberts, and Parish 2020). Nonetheless, studies using HPV16 containing cells suggest that the viral URR is hypomethylated upon cellular differentiation (Kim et al. 2003).

A recent study showed that papillomavirus virions package DNA with histones enriched in modifications typically associated with “active” (Porter et al. 2021). Of interest, the authors demonstrate that the levels of H3K4me3 were enriched on virions, compared to cellular controls Conversely, virions were depleted in H3K9me3 (Porter et al. 2021). There is emerging evidence of active associations between histone lysine methylation and DNA methylation (Rose and Klose 2014). For example, MeCP2 binds to methylated CpG (Nan et al. 1998), recruits the Suv39h1/2 histone methyltransferases (Fuks 2003), increasing H3K9me marks (Fuks 2003; Lunyak 2002). In parallel, the H3K9me mark recruits Dnmt3a/b to heterochromatin, leading to *de novo* methylation of CpG sites (Lehnertz et al. 2003; Otani et al. 2009). Since H3K9me3 is depleted in virions, it is tempting to conclude that virion DNA will be hypomethylated. Furthermore, H3K4me3 appears to be mutually exclusive with DNA methylation (M. Weber et al. 2007). H3K4me3 serves as a binding site for H3K9me2 demethylases (Horton et al. 2010), which would lead to loss of DNA methylation. Since virion DNA is enriched for H3K4me3, this further strengthens the hypothesis that viral DNA would be depleted in DNA methylation. Therefore, it seems reasonable to assume that infecting the virus genome will be hypomethylated and therefore serve as a TLR9 PAMP.

As mentioned, recognition by TLR9 would lead to an antiviral response. Indeed, siRNA-mediated knock-down of TLR9 has been shown to dramatically upregulate viral copy number and transcription following infection with HPV16 (Hasan et al. 2013), suggesting that TLR9 can restrict HPV infection. The observation that despite a reduction in viral CpG, HPV16 infection is still improved by interfering with TLR9 (signaling) demonstrates an important rule in host-pathogen interactions. While the loss of all (unmethylated) CpG dinucleotides would avoid detection by TLR9, the virus can likely not completely remove all CpGs from its genome. The virus and the host establish an uneasy balance.

### D3. Sustained flight and the bat immune system

Members of Chiroptera are classified into two suborders – Yinpterochiroptera (Rhinolophoid and megabats) and Yangochiroptera all other bat species (Lei and Dong 2016; Teeling et al. 2002; Springer et al. 2001). These suborders diverged roughly 60 million years ago (Lei and Dong 2016). Interestingly, the Yangochiroptera evolved flight and echolocation simultaneously, while the Yinpterochiroptera evolved these features separately (Anderson and Ruxton 2020). Sustained flight necessitated an increased metabolic capacity (Shen et al. 2010), which required bats to accommodate oxidative metabolism by-products such as DNA damage (Barzilai 2002). Indeed, many genes involved in DNA damage response and immunity have been demonstrated to be under positive evolutionary selection (Hawkins et al. 2019; Zhang et al. 2013). Since both suborders of bats ’invented’ flight independently, likely, the corresponding adaptations are also different. Indeed, we and others show that TLR9 is also under diversifying selection, specifically in the Yangochiroptera. While TLR9 likely did not evolve specifically to restrict papillomavirus infections, the virus likely needs to minimize its CpG content.

### D4. Direct evidence of co-evolution between the virus and its hosts

Co-evolution alongside their hosts has been suggested to be an essential factor in the evolution of papillomaviruses (Rector et al. 2007; Van Doorslaer 2013). However, the evolutionary history of PVs is complex. PVs isolated from fish form a monophyletic group distinct from those from mammals. However, within the mammalian papillomaviruses, there is no strict codivergence pattern that would unambiguously indicate an ancient relationship between host and virus. There has been no direct evidence in favor of co-evolution between papillomaviruses and their hosts. The co-evolution theory would predict that the virus would need to adapt when the host evolves a new skill. Therefore, as TLR9 evolves new functionalities, the virus would need to respond to preserve the balance between virus and host. Indeed, our data suggest that as Yangochiroptera TLR9 is undergoing diversifying selection, papillomavirus genomes infecting these bats further depleted their CpG content, specifically in the context of a known TLR9 PAMP. This is the first direct evidence of co-evolution between this family of viruses and their hosts.

In the phylogenetic analysis, the viruses that infect Yinpterochiroptera *and* Yangochiroptera, respectively, are not monophyletic but rather are present in three mixed clades (**Figure 2**; EsPV1, EsPV3, and RfPV1; MscP2 and EhPV1; TbraPV1-3). This suggests that these three main clades diverged before the ancestor of Yinpterochiroptera *and* Yangochiroptera split over 65 million years ago. As these ancestral viruses co-evolved with the Yangochiroptera hosts, they selected for loss of CpG. This occurred at least three separate times in the evolution of Yangochiroptera viruses. This strongly argues against a founder effect but in favor of recurring co-evolutionary interactions.

### D5. Immune evasion by nucleotide sequence editing

We previously used computer modeling and reconstruction of ancestral alphapapillomavirus genomes to show that these viruses depleted TpC depletion to allow for replication in tissues with high APOBEC3 expression – presumably to evade restriction by APOBEC3 by selecting for variants that contain reduced target sites in their genomes. We observed a similar correlation between TLR9 and CpG depletion, strengthening the notion that papillomaviruses avoid detection by the immune system by changing the nucleotide composition of their genomes without dramatically changing the protein-coding ability. This strategy likely allows the virus to maintain its core functionalities. Most viral proteins are multifunctional and interact with a plethora of host proteins. Amino acid level changes would likely disrupt these functions. The *de novo* evolution of new proteins is rare and is further complicated by the small genome size and overlapping open reading frames (Van Doorslaer and McBride 2016; Willemsen and Bravo 2019, 5).

### D6. Oncogene mediated reduction of TLR9

We propose that HPVs evade detection by TLR9 in the endosome by depleting CpG dinucleotides from their genomes. Interestingly, the E6 and E7 oncoproteins of different human papillomaviruses have been shown to downregulate the expression of TLR9 (Hasan et al. 2013; 2007; Pacini et al. 2015; 2017). Importantly, E6 and E7 are not delivered to the cell during infection but require onset of viral transcription after the viral genome is delivered to the nucleus and presumably has already been sensed in the endosome. This implies that the ability to degrade TLR9 may serve an additional function during the viral lifecycle, independent of initial infection. This idea is supported by the observation that other viruses (Merkel Cell Polyomavirus, Hepatitis B, and EBV) also interfere with TLR9 function during the maintenance phase of the infection (Fathallah et al. 2010; Vincent et al. 2011; Shahzad et al. 2013). Nonetheless, this oncogene-mediated repression of TLR9 occurs after infection and would still necessitate that the virus evades detection during infectious entry.

## D7. Conclusion

In conclusion, phylogenetic and genomic analyses of novel bat-associated viruses TbraPV2 and TbraPV3 demonstrate that host-virus interaction, specifically evasion of the innate immune system, affects the evolution of papillomaviruses. These data suggest that TLR9 acts as a restriction factor for papillomavirus infection. Furthermore, we provide the first direct evidence for co-evolution between papillomaviruses and their hosts.

## E. Materials and Methods

### E1. Data and Code availability

We retrieved full-length reference sequences from the PV database (PaVE; pave.niaid.nih.gov). Data and code for all analyses is available from https://github.com/KVDlab/King-2021. TbraPV2 (MW922427) and TbraPV3 (MW922428) sequences are available on GenBank. Raw sequencing data is available on SRA (PRJNA718335).

### E2. Sampling and sample processing

Bats were captured in mist nets set over water sources, extracted from the nets and put in brown paper bags. Bats were held in bags for 20 minutes, removed, and then measured and weighed. Individuals were identified to species in the field using metrics such as forearm length and weight. Feces and urine were collected from the bag or swabbed directly off the bat using a PurFlock 0.14" Ultrafine swab (Puritan, Guilford, Maine). All feces and urine samples were put into tubes containing 0.5 mL buffer consisting of 1x PBS and 50% Glycerol. These samples were held on ice until returning to the lab where they were stored in a -80°C freezer. All applicable international, national and institutional guidelines for the care and use of animals were followed during sampling. The study was approved by the University of Arizona Institutional Animal Care and Use Committee permit #15-583. Permits from the Arizona Department of Game and Fish were numbered SP506475.

Of each of the fecal samples, 5 g was homogenized in SM buffer and the homogenate was centrifuged at 6000 × g for 10 min. The supernatant was sequentially filtered through 0.45 μm and 0.2 μm syringe filters and viral particles in the filtrate were precipitated with 15% (w/v) PEG-8000 with overnight incubation at 4 °C followed by centrifugation at 10,000 ×g as described (Payne et al. 2020). The pellet was resuspended in 500μL of SM Buffer and 200μL of this was used for viral DNA extraction using the High Pure Viral Nucleic Acid Kit (Roche Diagnostics, Indianapolis, IN, USA). The total DNA was amplified using rolling circle amplification (RCA) with the TempliPhi 2000 kit (GE Healthcare, USA) and the RCA products used to prepare Illumina sequencing libraries then sequenced at Novagene Co. Ltd. (Hong Kong) on an Illumina NovaSeq 6000. The paired-end raw reads were trimmed using default settings within Trimmomatic v0.39 (Bolger, Lohse, and Usadel 2014) and the trimmed reads were *de novo* assembled using k-mer values of 33, 66, and 77 within metaSPAdes v 3.12.0 (Bankevich et al. 2012). The resulting contigs greater than 500 nucleotides were analyzed by BLASTx (Altschul et al. 1990) against a local viral protein database constructed from available NCBI RefSeq viral protein sequences (https://ftp.ncbi.nlm.nih.gov/refseq/release/viral/).

### E3. Calculation of nucleotide frequencies

To determine a single observed vs. expected (O/E) dinucleotide ratio across the entire viral genome, a custom python script was used that leverages the CompSeq program from Emboss (Warren, Van Doorslaer, et al. 2015). The expected frequencies of dinucleotide ‘words’ were estimated based on the observed frequency of single bases in the sequences. Only the forward frame was analyzed.

The tetramer content for each genome was calculated as described for the dinucleotides. To normalize tetramer content across groups of viruses we calculated the average O/E ratio across the different groups. These average O/E ratios were compared as indicated in the figure legends. To test whether any of the tested tetramers depletions are statistically significant, we randomly shuffled each viral genome. To ensure that each randomly shuffled sequence would maintain the same dinucleotide ratio as the original sequence, we used the Altschul and Erickson algorithm (Stephen F Altschul and Erickson Bruce W 1985) as implemented by Clote and colleagues (Clote 2005). Based on these shuffled sequences, we calculated the above Yangochiroptera/Yinpterochiroptera ratio. This was repeated 1000 times to establish a null distribution. The 1-percentile was used as a significance cutoff.

### E4. Phylogenetic analyses

Annotated sequences (n = 409) were downloaded from the PaVE genome database. A maximum likelihood phylogenetic tree was constructed as described (King and Van Doorslaer 2018). The amino-acid sequences for E1, E2, and L1 of all known papillomaviruses and the new TbraPV2 and TbraPV3 were individually aligned in MAFFT v7.3 (Katoh 2002; 2005) using the L-INS-I algorithm. A partitioning scheme for the concatenated E1-E2-L1 alignment was determined under corrected Akaike information criterion (AICc) implemented in PartitionFInder2 (Lanfear et al. 2017), which separately identified each gene to evolve under the LG+I+G+F evolutionary substitution model. The concatenated E1-E2-L1 alignment was used to infer the best maximum likelihood (ML) phylogenetic tree using RAxML-HPC v.8 (Stamatakis 2014) on CIPRES science gateway (Miller, Pfeiffer, and Schwartz 2010) followed by a rapid bootstrapping analysis. A posteriori bootstopping was automatically rendered in RAxML under the extended majority-rule consensus tree criterion (autoMRE). The best ML tree was rendered and edited in RStudio using the ‘ggtree’ (Yu et al. 2018) and ‘treeio’ (L.-G. Wang et al. 2020) packages.

Taxonomic classification of TbraPV2 and TbraPV3 was based on pairwise sequence identity. The L1 sequence of each pair was aligned at the amino acid level using the L-INS-I algorithm as implemented within the MAFFT v7.3 (Katoh 2002; 2005). This way the alignments preserve the codons. The resulting alignments are back translated to nucleotide alignments and used to calculate pairwise sequence identity.

### E5. Coevolution analysis

We used functions in the R ‘ape’ (Paradis and Schliep 2019) package to extract a well-supported clade from the maximum likelihood phylogenetic tree. The extracted clade represents contains viral sequences in the genera *Lambdapapillomavirus*, *Mupapillomavirus*, *Nupapillomavirus*, *Kappapapillomavirus*, *Sigmapapillomavirus*, and *Dyosigmapapillomavirus*, and the largest set of known of bat papillomaviruses, including the two novel bat papillomaviruses described in this paper. A corresponding host species phylogeny was downloaded from TimeTree (www.timetree.org) (Hedges, Dudley, and Kumar 2006; Kumar et al. 2017; Hedges et al. 2015). A tanglegram representing the evolutionary relationship between the papillomaviruses and their hosts was constructed in the ‘phytools’ package (Revell 2012). Phytools will optimize the tanglegram by rotating nodes in the rooted phylogenies to minimize crossings between connecting lines between both trees.

An additional subtree was extracted to minimize the impact of the genus *Lamdapapillomavirus*. The viral types included in this smaller dataset are underlined in **Figure 2**. To assess the congruency between PV and host phylogenies, we used the Procrustes Approach to Cophylogenetic Analysis (PACo) (Balbuena, Míguez-Lozano, and Blasco-Costa 2013) as implemented in R for both datasets. Briefly, PACo uses cophenetic distance matrices for the virus and host trees and an association matrix of virus -host interactions. To assess statistical significance, a Procrustean super-imposition of the sum of squared residuals was generated from 1000 network randomizations under the “r2” randomization model. Under this model, host specialization is assumed to drive the virus diversification (Hutchinson et al. 2017). The values for the actual tree comparisons were considered statistically significant if they fell outside the 95% confidence interval (C.I.)

To quantify the similarity between the virus and host phylogenies, we calculated the Wasserstein distance using the ‘castor’ R package (Louca and Doebeli 2018). The Wasserstein distance is based on a modified graph Laplacian (MGL). The MGL uses evolutionary distances between nodes to construct a matrix which maintains branch length and tree topology information and allows for the comparison of phylogenies from different species. Specifically, the differences between a phylogeny’s degree matrix (sum of branch lengths from one node n to all others) and distance matrix (sum of all pairwise branch lengths) is calculated to generate a spectrum of eigenvalues. To calculate a normalized MGL (nMGL) the MGL is divided by the degree matrix. The normalized MGL is specifically useful when comparing trees on different timescales by emphasizing topology over size (Lewitus and Morlon 2016). The Wasserstein distance represents the largest eigenvalue from the spectra of the modified graph Laplacians. All eigenvalues from the graph Laplacian spectrum were used to calculate the Wasserstein tree distance. The Wasserstein tree distance metric calculated in ‘castor’ considers branch length and tree topology, takes values between 0 and 1. Identical tree topologies would have a Wasserstein distance of 0.

### E6. Analysis of codon usage

A custom script was used to delete all overlaps between open reading frames. Briefly, overlaps between E6 and E7, E1 and E8, E2 and E4, L2 and L1 were removed when present. For each overlap, entire codons were removed as not introduce frameshifts. These sequences were concatenated and further analyzed.

Cusp (Emboss suite of tools) was used to generate codon usage tables for each virus. These tables were compared using codcmp (Emboss suite of tools). For each codon in the table codcmp calculates the proportion of a codon to the total number of the codons in the. Next, codcmp calculates the difference between the usage fractions in both tables.

The amino acid composition for each sequence was calculated as described (Carugo 2008).

### E7. Diversifying selection of analysis of Yangochiroptera TLR9

TLR9 sequences were downloaded from NCBI and translated into putative proteins. The amino acid sequences were aligned using MAFFT v7.3, and back translated into codon-aware nucleotide alignments. FastTree (Price, Dehal, and Arkin 2010) was used to construct a maximum likelihood phylogenetic tree using the GTR substitution model of evolution.

To determine whether the strength of natural selection intensified along the along Yangochiroptera compared to the Yinpterochiroptera, we used RELAX. After fitting a codon model with three ω classes to the phylogeny (null model), RELAX then tests for changes to the intensity of selection by introducing a selection parameter k. The null and alternative models are compared using a Likelihood Ratio Test. A significant result of k>1 indicates that selection strength has been intensified along the test branches (Wertheim et al. 2015).

aBSREL (adaptive Branch-Site Random Effects Likelihood) was used to test if positive selection has occurred on the branches leading to Yangochiropter*a*. aBSREL determines whether a proportion of sites have evolved under positive selection (Smith et al. 2015).

Finally, FEL (Fixed Effects Likelihood) was used to infer non-synonymous (dN) and synonymous (dS) substitution rates on a per-site basis. This method assumes that the selection pressure for each site is constant along the entire phylogeny. FEL fits a MG94xREV model to each codon site to infer nonsynonymous and synonymous substitution rates at each site. A Likelihood Ratio Test determines if dN is significantly greater than dS.

### E8. Statistical analysis

One- or two-way analysis of variance (ANOVA) were used where appropriate. Data are presented as box-and-whisker plots with Tukey’s method for outliers noted as distinct data points. All graphs were generated using R. Results were considered statistically significant at a *P*-value of < 0.05.

